## Supplementary Data for "Oncogenic drivers dictate immune control of acute myeloid leukemia"

#### Supplementary Material and Methods

### Oncogenic drivers dictate immunological control of disease progression in acute myeloid leukemia.

#### Supplementary Figure legends

**Supplementary Figure 1:** (A) Number of CFU-GM colonies grown in methylcellulose from wild type (WT) and Rag2$^{-/-}$γc$^{-/-}$ mice. (B) Kaplan-Meier curves for Rag2$^{-/-}$γc$^{-/-}$ primary recipients transplanted with Rag2$^{-/-}$γc$^{-/-}$ HSPCs subsequent to retroviral transduction with constructs expressing the oncogenes BCR-ABL and NUP98-HOXA9 (BA/NH), MLL-AF9 (MA9) or AML1-ETO and Nras$^{G12D}$ (AE/Nras$^{G12D}$) (top). Cell morphology of secondary BA/NH, MA9 and AE/Nras$^{G12D}$ AML in H&E stained spleens of Rag2$^{-/-}$γc$^{-/-}$ recipients (middle). Cell surface expression by flow cytometry of CD11b and cKIT on GFP+ primary BA/NH, MA9 and AE/Nras$^{G12D}$ AMLs passaged through Rag2$^{-/-}$γc$^{-/-}$ recipients (bottom). (C) PCR using gDNA isolated from GFP+ AML cells from BA/NH and AE/ Nras$^{G12D}$ primary recipients and primers spanning the AML1-ETO, BCR-ABL and NUP98-HOXA9 breakpoints and exon 4 and 5 in Nras. Amplification of GAPDH was used as a positive control. Spleen weights of secondary recipients transplanted with primary (D) BA/NH, (E) MA9, and (F) AE/Nras$^{G12D}$. (G) Percentage of GFP+ 2$^{o}$ MA9 AML cells in BM 24h post transplant. (H) Kaplan-Meier curves comparing survival between Rag2$^{-/-}$γc$^{-/-}$ (n=4) and WT (n=5) secondary recipients transplanted with 1,000 1$^{o}$ BA/NH AML cells. Each point represents a biological replicate. Mann-Whitney test for comparison between two groups. Mantel-Cox test for comparison of Kaplan-Meier curves, *p <.05, ** p < 0.01.

**Supplementary Figure 2:** (A) Gating strategy for monocytes/granulocytes (CD11b+). (B) Histograms comparing expression of $H2D^b$, $H2K^b$ and MHC Class II (IA/E) on CD11b+ cells from naïve mice and on the cell surface of GFP+ BA/NH, MA9 and $Nras^{G12D}$ cells. (C) PCR using gDNA isolated from GFP+ AML cells from BA/NH and AE/$Nras^{G12D}$ primary and secondary recipients and primers spanning the AML1-ETO breakpoint and exon 4 and 5 in Nras. Amplification of GAPDH was used as a positive control. (D) Median fluorescence intensity (MFI) of MHC Class II (IA/E) on cell surface of CD11b+ GFP+ cultured hematopoietic stem and progenitor cells (HSPCs) isolated from $Rag2^{-/-}\gamma c^{-/-}$ bone marrow, 72hrs post transduction with individual retroviral constructs expressing the oncogenes listed on the x-axis or an empty vector (EV) control. MFI expressed as fold change EV control, n=3 independent HPSC donors and transductions. (E) Microarray derived gene expression of HLA-DQB1, HLA-DOA, HLA-DMA, HLA-DPA1 and HLA-DPB1 from bulk PB/BM of n=10 MLL-X and 42 mutant NRAS AML patients[1]. One-way ANOVA with Tukey's multiple testing correction (D), Mann-Whitney test for pairwise comparisons between groups (E).

**Supplementary Figure 3:** (A) Histograms comparing expression of CD80, CD86, PD-L1, GAL-9 and CD155 on CD11b+ cells from naïve mice and on the cell surface of GFP+ BA/NH, MA9 and $Nras^{G12D}$ cells. (B) Median fluorescence intensity (MFI) of CD80, CD86, PD-L1, GAL-9 and CD155 on cell surface of CD11b+ GFP+ cultured hematopoietic stem and progenitor cells (HSPCs) isolated from $Rag2^{-/-}\gamma c^{-/-}$ bone marrow, 72hrs post transduction with individual retroviral constructs expressing the oncogenes listed on the x-axis or an empty vector (EV) control. MFI expressed as fold change EV control, n=3 independent HPSC donors and transductions. One-way

ANOVA with Tukey's multiple testing correction (B: CD86, GAL-9, CD155), Kruskal-Wallis test (B: CD80, PD-L1). * p <0.05, ** p < 0.01, *** p <0.001, **** p <0.0001.

**Supplementary Figure 4:** (A) Frequency of T cells (GFP-/CD3+/NK1.1) as a percentage of GFP- mononuclear cells in the peripheral blood of wild type mice treated with T cell depleting antibodies (anti-CD4/anti-CD8) or isotype control and transplanted with MA9 AML, as expressed as a percentage of the average for the isotype control per timepoint. (B) Gating strategy and representative flow plots for naïve (CD44- CD62L+), T effector memory (TEM, CD44+ CD62L-) and central memory (TCM, CD44+ CD62L+) CD4+ and CD8+ T cells from naïve mice and BA/NH, MA9 and $Nras^{G12D}$ AML recipients. (C) Frequency of naïve and TCM CD4+ T cells and (D) CD8+ T cells in the spleens of naïve wild type C57BL/6J mice and BA/NH, MA9 and $Nras^{G12D}$ AML recipients. One way ANOVA with Tukey's p-value adjustment. * p <0.05, ** p < 0.01, *** p <0.001, **** p <0.0001.

**Supplementary Figure 5:** (A) Representative histograms displaying expression of PD-1, DNAM-1, TIM-3, and KLRG1 on CD4+ T cells from BA/NH, MA9 and $Nras^{G12D}$ immunocompetent recipients. (B) Representative flow plots of CD4+ T cells from BA/NH, MA9 and $Nras^{G12D}$ immunocompetent recipients showing co-expression of PD-1 and DNAM-1, PD-1 and TIM-3, and PD-1 and KLRG1. (C) Representative histograms displaying expression of PD-1, DNAM-1, TIM-3 and KLRG1 on CD8+ T cells from BA/NH, MA9 and $Nras^{G12D}$ immunocompetent recipients. (D) Representative flow plots of CD8+ T cells from BA/NH, MA9 and $Nras^{G12D}$ immunocompetent recipients showing co-expression of PD-1 and DNAM-1, PD-1 and TIM-3, and PD-1 and KLRG1.

**Supplementary Figure 6:** (A) Experimental schema for the generation of immunoedited (IE) and non-immunoedited (N-IE) $Nras^{G12D}$ cells for RNA sequencing. (B) Principal component analysis of RNA sequencing data generated from GFP+ $Nras^{G12D}$ AML cells isolated from either immunocompetent WT (IE, n=5) or immunodeficient $Rag2^{-/-}\gamma c^{-/-}$ (N-IE, n=5) recipients. (C) Quantification of Nras normalized read-count in N-IE and IE $Nras^{G12D}$ cells. (D) Quantification of Nras gene expression by qPCR in N-IE and IE $Nras^{G12D}$ cells. (E) Quantification of Ki67 immunohistochemistry on tumour-bearing regions of the spleen and liver isolated from either $Rag2^{-/-}\gamma c^{-/-}$ or WT secondary recipients of $Nras^{G12D}$ AML. (n=4 independent recipients, each data point represents the mean of 5 separate fields). (F) Quantification of phosphorylated histone H3 (p-H3) immunohistochemistry on tumour-bearing regions of the spleen and liver isolated from either $Rag2^{-/-}\gamma c^{-/-}$ or WT secondary recipients of $Nras^{G12D}$ AML. (n=4 independent recipients, each data point represents the mean of 5 separate fields). (G) Lack of enrichment of genes correlating with down-regulation of Nras signaling and upregulation of Myc targets in MA9 AML passaged in WT (n=4) compared to $Rag2^{-/-}\gamma c^{-/-}$ (n=5) recipients, as determined from RNA-sequencing of GFP+ AML cells. (H) Enrichment of genes correlating with an interferon gamma response in MA9 AML passaged in WT (n=4) compared to $Rag2^{-/-}\gamma c^{-/-}$ (n=5) recipients, as determined from RNA-sequencing of GFP+ AML cells. (I) Enrichment of genes correlating with an interferon gamma response in non-immunoedited (N-IE) $Nras^{G12D}$ AML, as determined from RNA-sequencing of GFP+ AML cells isolated from either immunocompetent WT (IE, n=5) or immunodeficient $Rag2^{-/-}\gamma c^{-/-}$ (N-IE, n=5) recipients. Mann-Whitney test for comparison between two groups (C, D, E, F).

#### Supplementary methods

##### Colony forming assay

BM from 4 x Rag2-/-γc-/- and 4 x wild type C57BL/6J was harvested and red blood cells lysed. 10,000 BM cells per mouse were added to 3mL of MethoCultTM GF M3434 (STEMCELL Technologies), mixed thoroughly by vortexing prior to dispensing 1mL aliquots in triplicate into 35mm plates. Plated cells were incubated (37$^{o}$C, 5% $CO_2$) for 7 days and colony forming units-granulocyte, macrophage (CFU-GM) were visually enumerated.

##### Generation of murine AML models

Plasmids pMSCV-MLL-AF9-IRES-GFP (pMIG-MA9), pMSCV-NUP98-HOXA9-IRES-GFP (pMIG-NH9), pMSCV-BCR-ABL-IRES-GFP (pMIG-BA), pMSCV-AML1-ETO-IRES-GFP (pMIG-AE), pMSCV-IRES-GFP and the packaging plasmid pCL-Eco were a gift from Dr. D.G. Gilliland (Boston, MA). pMSCV-GFP-IRES-Nras$^{G12D}$ (pMIG-Nras$^{G12D}$) was a gift from A/Prof Ross Dickins (Melbourne, Australia). pMSCV-Myc-IRES-mCherry was a gift from Dr Gretchen Poortinga (Melbourne, Australia). Sanger sequencing was used to determine the exact sequence over the breakpoint of each fusion protein encoded in the listed constructs, using the primers detailed in the table below:

| Fusion Oncogene | Breakpoint | Sequencing Primer | Sequencing product |
|---|---|---|---|

| | | | |
|---|---|---|---|
| AML1-ETO (RUNX1-RUNX1T1) | Exon6 – Exon3 | AML-ES 5’GAGGGAAAAGCTTCACTCTG3’ | CCTACCACAGAGCCATCAAAATCACAGTGGATGGGCCCCGAGAACCTCGAAATCGTACTGAGAAGCACTCCACAATGCCAGACTCACCTGTGGATGTGAAGACGCAATCTAGGCTGACTCCTCCAACAATGCCACCTCCCCCAACTACTCAAGGAGCTCCAAGAACCAGTTCATTTACACCGACAACGTTAACTAATGGCACGAGCCATTCTCCTACAGCCTTGAATGGCGCCCCCTCACCACCCAATGGCTTCAGCAATGGGCCTTCCTCTTCTTCCTCCTCTCTGGCTAATCAACAGCTGCCCCCAGCCTGTGGTGCCAGGCAACTCAGCAAGCTGAAAAGGTTCCTTACTACCCTGCAGCAGTTTGGCAATGACATTTCACCCGAGATAGGAGAAAGAGTTCGCACCCTCGTTCTGGGACTAGTGAACTCCACTTTGACAATTGAAGAATTTCATTCCAAACTGCAAGAAGCTACTAACTTCCCACTGAGACCTTTTGTCATCCCATTTTTGAAGGCCAACTTGCCCCTGCTGCAGCGTGAGCTCCTCCACTGCGCAAGACTG |
| MLL-AF9 (KMT2A-MLLT3) | Exon10 – Exon9 | MLL-F1 5’CGCCTCAGCCACCTACTACAG3’ | TCCTAGTGAGCCCAAGAAAAAGCAGCCTCCACCACCAGAATCAGGTCCAGAGCAGAGCAAACAGAAAAAAGTGGCTCCCCGCCCAAGTATCCCTGTAAAACAAAAACCAAAAGAAAAGGAAAAACCACCTCCGGTCAATAAGCAGGAGAATGCAGGCACTTTGAACATCCTCAGCACTCTCTCCAATGGCAATAGTTCTAAGCAAAAAATTCCAGCAGATGGAGTCCACAGGATCAGAGTGGACTTTAAGGAAGACTGTGAAGCAGAAAATGTGTGGGAGATGGGAGGCTTAGGGATCCTTGAAGTGAAAAGTCCAATAAAGCAAAGCAAATCAGATAAGCAAATAAAGAATGGTGAATGTGACAAGGCATACCTAGATGAACTGGTAGAGCTTCACAGAAGGTTAATGACATTGAGAGAAAGACACATTCTGCAGCAGATCGTGAACCTTATAGAAGAAACTGGACACTTTCATATCACAAACACAACATTTGATTTTGATCTTTGCTCGCTGGACAAAACCACAGTCCGTAAACTACAGAGTTACCTGGAAACATCTGGAACATCCTGAG |
| BCR-ABL1 | Exon14-Exon2 | BCRP210-B2-C 5’ CAGATGCTGACCAACTCGTGT3’ | GATGATGAGTCTCCGGGGCTCTATGGGTTTCTGAATGTCATCGTCCACTCAGCCACTGGATTTAAGCAGAGTTCAAAAGCCCTTCAGCGGCCAGTAGCATCTGACTTTGAGCCTCAGGGTCTGAGTGAAGCCGCTCGTTGGAACTCCAAGGAAAACCTTCTCGCTGGACCCAGTGAAAATGACCCCAACCTTTTCGTTGCACTGTATGATTTTGTGGCCAGTGGAGATAACACTCTAAGCATAACTAAAGGTGAAAAGCTCCGGGTCTTAGGCTATAATCACAATG |
| NUP98-HOXA9 | Exon12 – Exon1 | NUP98 forward-3 5’GCACAAATACCAGTGGGAATA 3’ | GGACTGGGCTTGGTGCAGGATTTGGAACAGCTCTTGGTGCTGGACAGGCATCTTTGTTTGGGAACAACCAACCTAAGATTGGAGGGCCTCTTGGTACAGGAGCCTTTGGGGCCCCTGGATTTAATACTACGACAGCCACTTTGGGCTTTGGAGCCCCCCAGGCCCCAGTAGTTGATAGAGAAAAACAACCCAGCGAAGGCGCCTTCTCTGAAAACAATGCTGAGAATGAGAGCGGCGGAGACAAGCCCCCCATCGATCCCAATAACCCAGCAGCCAACTGGCTTCATGCGCGCTCCACTCGGAAAAAGCGGTGCCCCTATACAAAACACCAGACCCTGGAACTGGAGAAAGAGTTTCTGTTCAACATGTACCTCACCAGGGACCGCAGGTACGAGGTGGCTCGACTGCTCAACCTCACCGAGAGGCAGGTCAAGATCTGGTTCCAGAACCGCAGGATGAAAATGAAGAAAATCAACAAAGACCGAGCAAAAGACGAGTGATGCCATTTGGGCTTATTTAGA |

Plasmid DNA was isolated from transformed E.coli using PureLink™ HiPure Plasmid Filter Maxiprep Kit (ThermoFisher) according to manufacturer’s instructions, with specific oncogenic sequences confirmed using  Sanger sequencing. Retroviruses were packaged by co-transfection of plasmids using FuGENE (Roche) into HEK293T cells.

Bone marrow (BM) from *Rag2*$^{-/-}$*γc*$^{-/-}$ mice (8-12 weeks of age) was depleted using Ter119, Gr1 and CD11b biotin conjugated antibodies and Dynabeads (Invitrogen) according to the manufacturer's instructions. Lineage-depleted BM cells at 2 x $10^{\wedge 6}$ cells/mL were cultured overnight (37°C, 5% $CO_2$) in RPMI supplemented with 10% FCS, 100 IU/mL Pencillin/Streptomycin, 10 ng/mL murine recombinant IL-3 (Peprotech), murine recombinant IL-6 (Peprotech) and 50 ng/mL murine recombinant stem cell factor (SCF) (Peprotech). Cells were re-suspended with 1 mL of unconcentrated retrovirus per well with polybrene (8 μg/mL) and HEPES (30 μL/mL) and mIL-3 10ng/ml, mIL-6 10ng/ml and mSCF 50ng/ml (Peprotech). Transduction combinations included: MLL-AF9 alone; BCR-ABL and NUP98-HOXA9; or AML1-ETO and Nras$^{G12D}$ expressed using the retroviral backbone MSCV-IRES-GFP.

BM cells were transduced by two room temperature spin infections for 90 mins at 3000 rpm separated by three-hour incubation at 37°C, 5% $CO_2$. For primary (1°) transplants, BM cells were assessed for viability and GFP expression by flow cytometry prior to transplantation by lateral tail vein injection into primary Rag2$^{-/-}$γc$^{-/-}$ recipients. For secondary (2°) and tertiary (3°) transplants, equal numbers of GFP$^{+}$ AML cells were transplanted into non-irradiated wild type C57BL/6J or Rag2$^{-/-}$γc$^{-/-}$ recipients (150K per recipient).

Cryopreserved 1° NrasG12D AML-bearing splenocytes were recovered and immediately transduced with either pMSCV-Myc-IRES-mCherry or the empty vector backbone retrovirus in short-term culture, as detailed above. Double positive GFP+, mCherry+ cells were sorted (BD FACSAria™) and expanded via transplantation in Rag2$^{-/-}$γc$^{-/-}$ recipients.

**Genotyping of murine AML models**

Genomic DNA was isolated from sorted GFP+ cells using QuickExtract™ DNA Extraction Solution (Illumina) according to the manufacturer's instruction. PCR was performed using the following primers: NUP98-HOXA9 (Fwd-5'gcacaaataccagtgggaata 3', Rev-5'gggcaccgctttttccgagtg 3', 373bp product), BCR-ABL (Fwd-5'cagatgctgaccaactcgtgt 3', Rev-5'gtttgggcttcacaccattcc 3', 377bp product), AML1-ETO (Fwd-5'gagggaaaagcttcactctg 3', Rev-5'gaaggcccattgctgaagc 3', 325bp product), NRas$^{G12D}$ ORF (Fwd-5'ccagtacatgaggacaggcg 3', Rev-5'acttgttgcctaccagcacc 3', 145bp product)(Fwd-5'ggacacagctggacaagagg 3', Rev-5'cacacttgttgcctaccagc 3', 190bp product).

**Histology**

Histology samples were processed by the QIMR Berghofer Histology Facility. Briefly tissues were fixed in 10% neutral buffered formalin, embedded in paraffin prior to staining with haematoxylin Aarmstadt Hx Crystals (Merck) and Eosin Y (H&E). In addition, tissues were stained for peroxidase labeled Ki67 (SP6, ab16667, Abcam) and phosphorylated histone H3 (p-H3) (polyclonal, 06-570, Merck Millipore) with images captured on a Nikon Eclipse Ci, DS-Fi2 microscope.

**RNA sequencing and bioinformatics analysis**

Oligo d(T) captured mRNA (100 μg) was processed for Next Generation Sequencing (NGS) using the NEB Next Ultra II RNA Library Prep Kit for Illumina (New England Biolabs). Quality was assessed using the High Sensitivity DNA Kit (Agilent) on the Agilent 2100 Bioanalyser with quantification using the Qubit DNA HS Assay Kit (Molecular Probes). Final libraries were sequenced using a high output single -end 75bp flow cell (version 2) on the Illumina Nextseq 550 platform. Reads were trimmed for adapter sequences using Cutadapt (version 1.11) and aligned using STAR [2] (version 2.5.2a) to the GRCm38, with assembly using the gene, transcript, and exon features of Ensembl (release 67). Expression was estimated using RSEM (version 1.2.30), with transcripts with zero read counts across all samples removed prior to analysis. Normalisation of read counts was performed by dividing by million reads mapped to generate counts per million (CPM), followed by the trimmed mean of M-values (TMM) method from the edgeR package [3]. For the differential expression analysis, reads were filtered but not normalized, since edgeR performs normalisation (library size and RNA composition) internally. The glmFit function was used to fit a negative binomial generalised log-linear model to the read counts for each transcript. Transcript wise likelihood ratio tests were conducted for each comparison. Log2 transformed, normalized read counts were used for heatmaps and principle component analysis (PCA).

Gene set enrichment analysis (GSEA) was performed using GSEA from Broad Institute [4, 5]. P-values were generated form 1000 gene set permutations, excluding gene sets with more than 3000 genes or less than 5 genes against custom made gene sets and Broads Hallmark and C2 database.

**Quantitative PCR analysis**

Primers used: Nras: Forward 5’ TGTTGGGAAAAGCGCCTTGA 3’ and reverse 5’ CCTGTCCTCATGTACTGGTCT 3’. Beta-actin (Actβ): Forward 5’ GACGATATCGCTGCGCTGGT 3’ and reverse 5’ CCACGATGGAGGGGAATA 3’. Primer specificity was determined by melt-curve analysis. Gene expression and copy number were quantified by Sybr green reaction (Life technologies) using ABI Viia7 qPCR machine according to standard protocols. Gene expression was determined by standard curve analysis and normalized to Actβ with relative gene expression and copy number expressed as a fold change.

**Publicly available microarray and single cell RNA-sequencing data analysis**

Microarray array data with accession GSE6981 were downloaded, pre-processed and normalized as previously described[1]. Samples from AML patients with NRAS mutation (n=42) and MLL-translocation (n=10) were extracted for analysis. Using the GSVA R package function ssgsea the relative enrichment of gene sets across was assessed for all genetic aberrations with more than 3 patients for genes encoding MHC Class II and for NRAS mutant samples was assessed for gene sets associated with cytolytic immune infiltration in AML [6] or Myc gene target activation [7].

Pre-processed and annotated single cell RNA-sequencing data of 6 AML patients with MLL-translocation, 6 patient with RAS mutations (2 KRAS,3 NRAS) and 7 healthy controls were obtained from GEO with accession GSE185381 [8]. For MHC II gene expression comparison across oncogenes, CD33 expressing malignant annotated cells were extracted and B, T and NK cells excluded. For each sample

MHC II gene expression was averaged.  For comparison of the percentage of PDCD1 expressing cells oncogenes annotated T cells were extracted.

**Supplementary Table 1:** Flow cytometry antibodies

| Marker | Clone | Manufacturer |
|---|---|---|
| TCR-β | H57-597 | Biolegend |
| CD4 | RM4-5 | Biolegend |
| CD8 | 53-6.7 | Biolegend |
| H-2D$^{b}$ | 28-14-8 | BD Biosciences |
| H2-K$^{b}$ | AF6-88-5 | BD Biosciences |
| CD80 | 16-10A1 | eBioscience |
| CD86 | GL-1 | Biolegend |
| CD155 | 4.24.3 | Biolegend |
| PD-L1 | 10F.9G2 | Biolegend |
| TIM-3 | RMT3-23 | eBioscience |
| MHC Class II | M5/114.15.2 | eBioscience |
| GAL-9 | 108A.2 | Biolegend |
| CD44 | IM7 | Biolegend |
| CD62L | MEL-14 | Biolegend |
| PD-1 | RMPI-30 | Biolegend |
| CD3 | 17A2 | Biolegend |

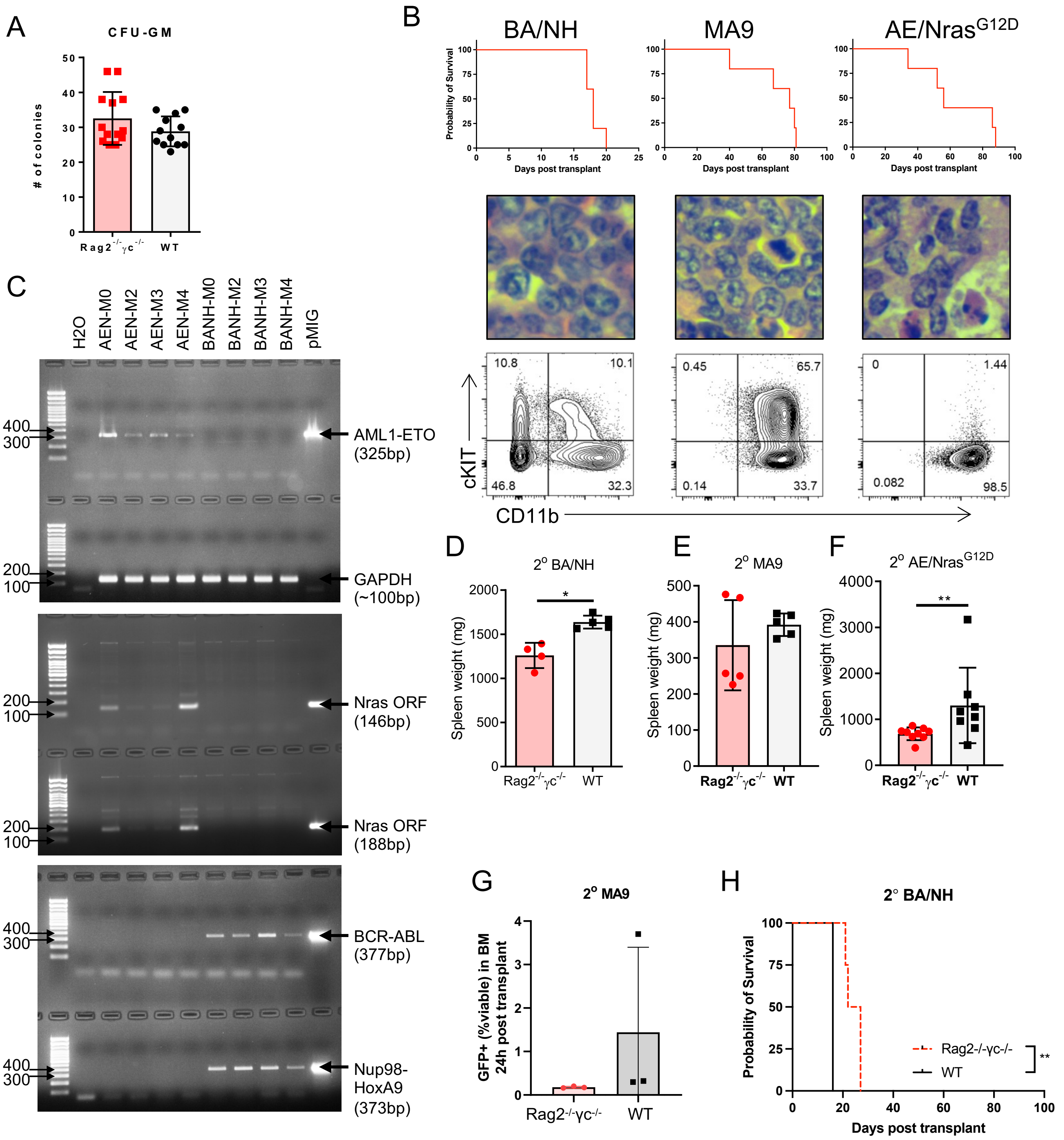


Supplementary Figure 1

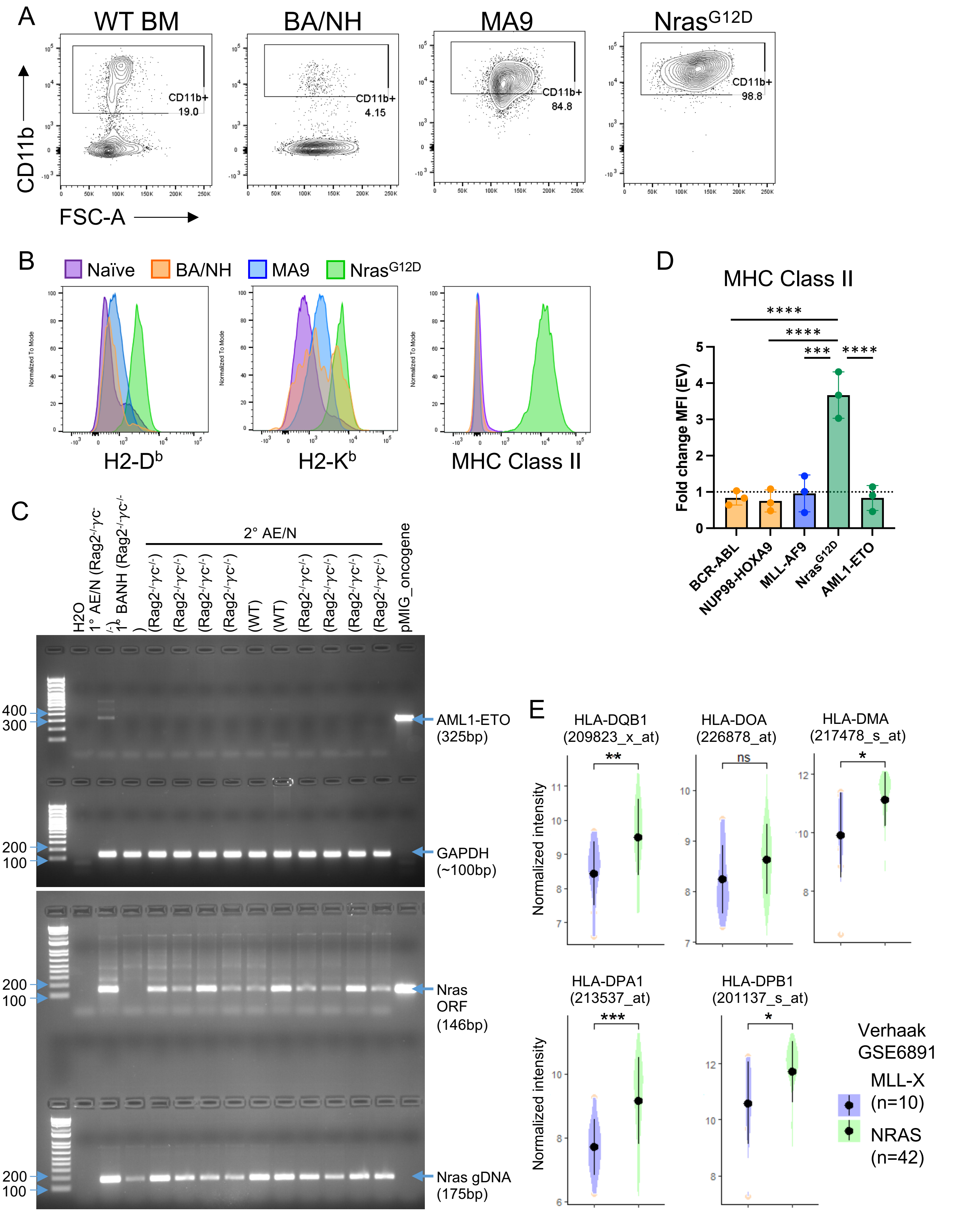


Supplementary Figure 2

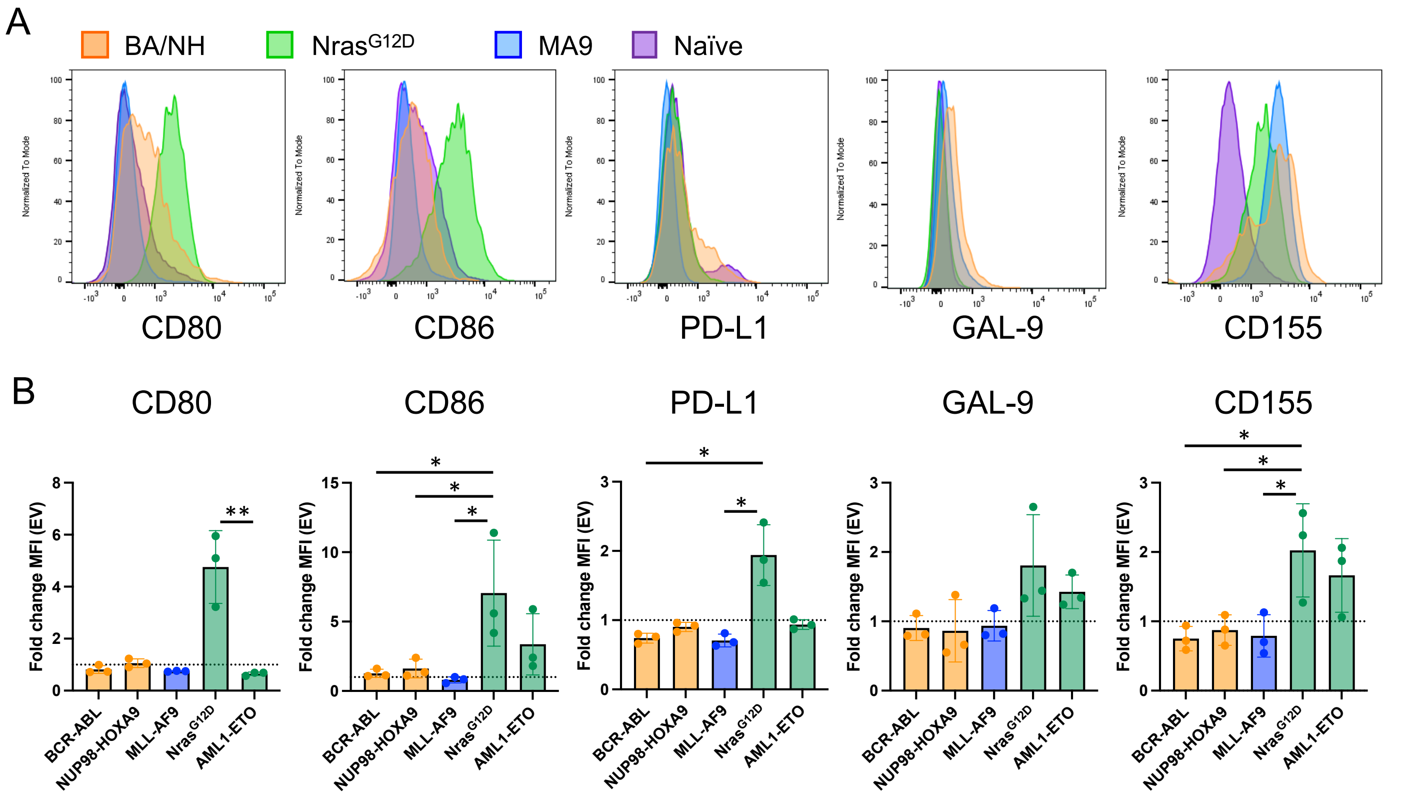


Supplementary Figure 3

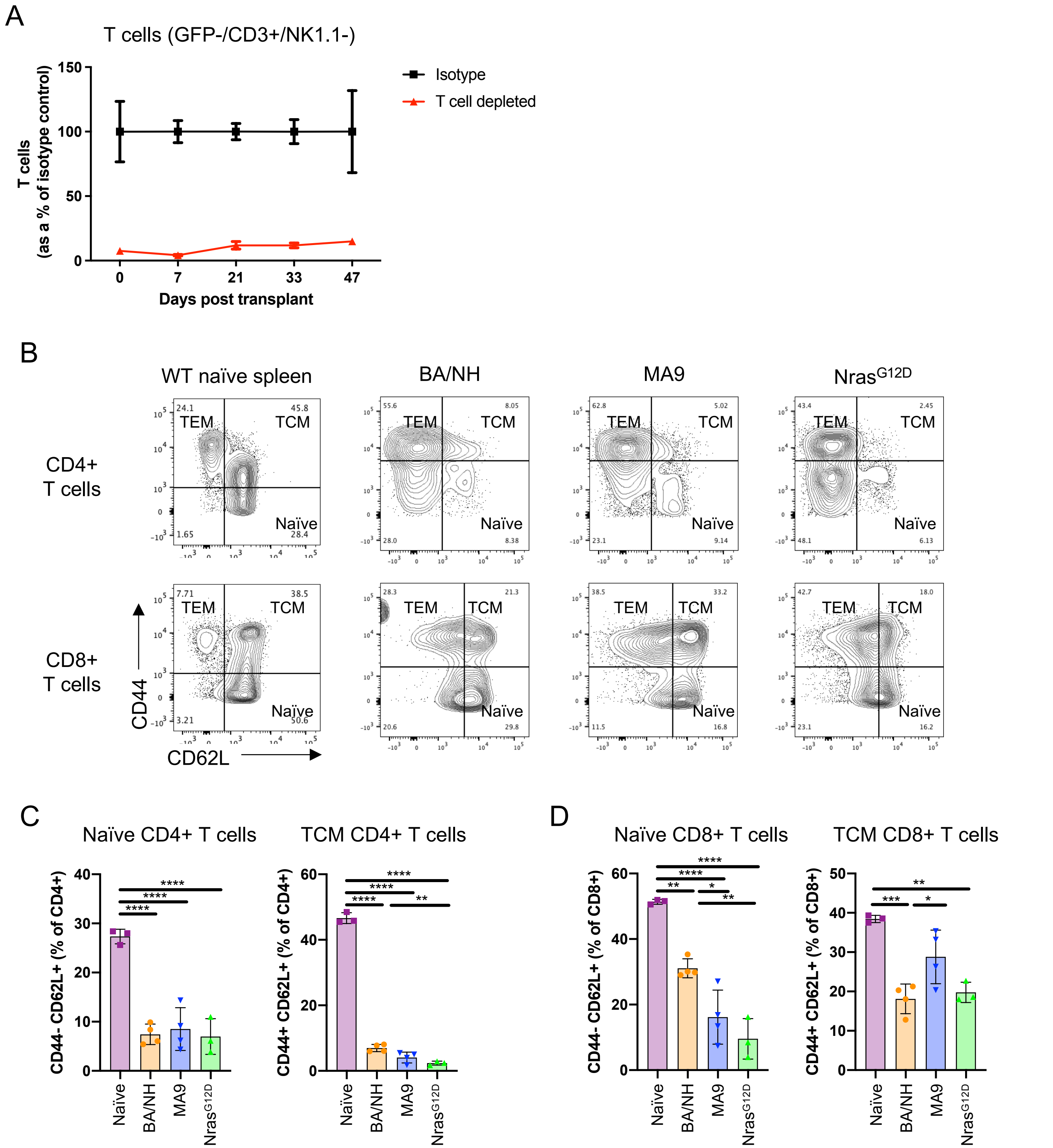


Supplementary Figure 4

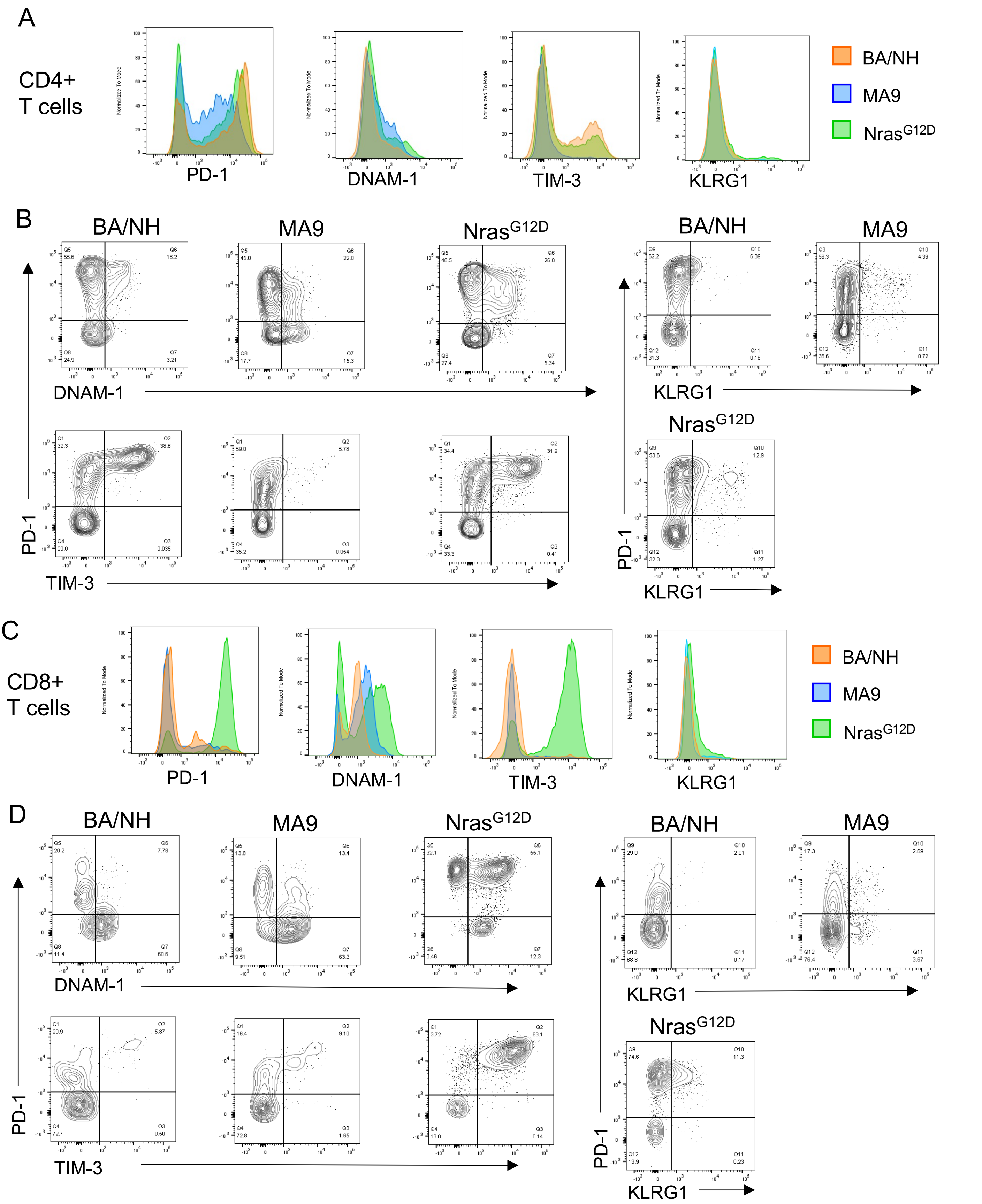


Supplementary Figure 5

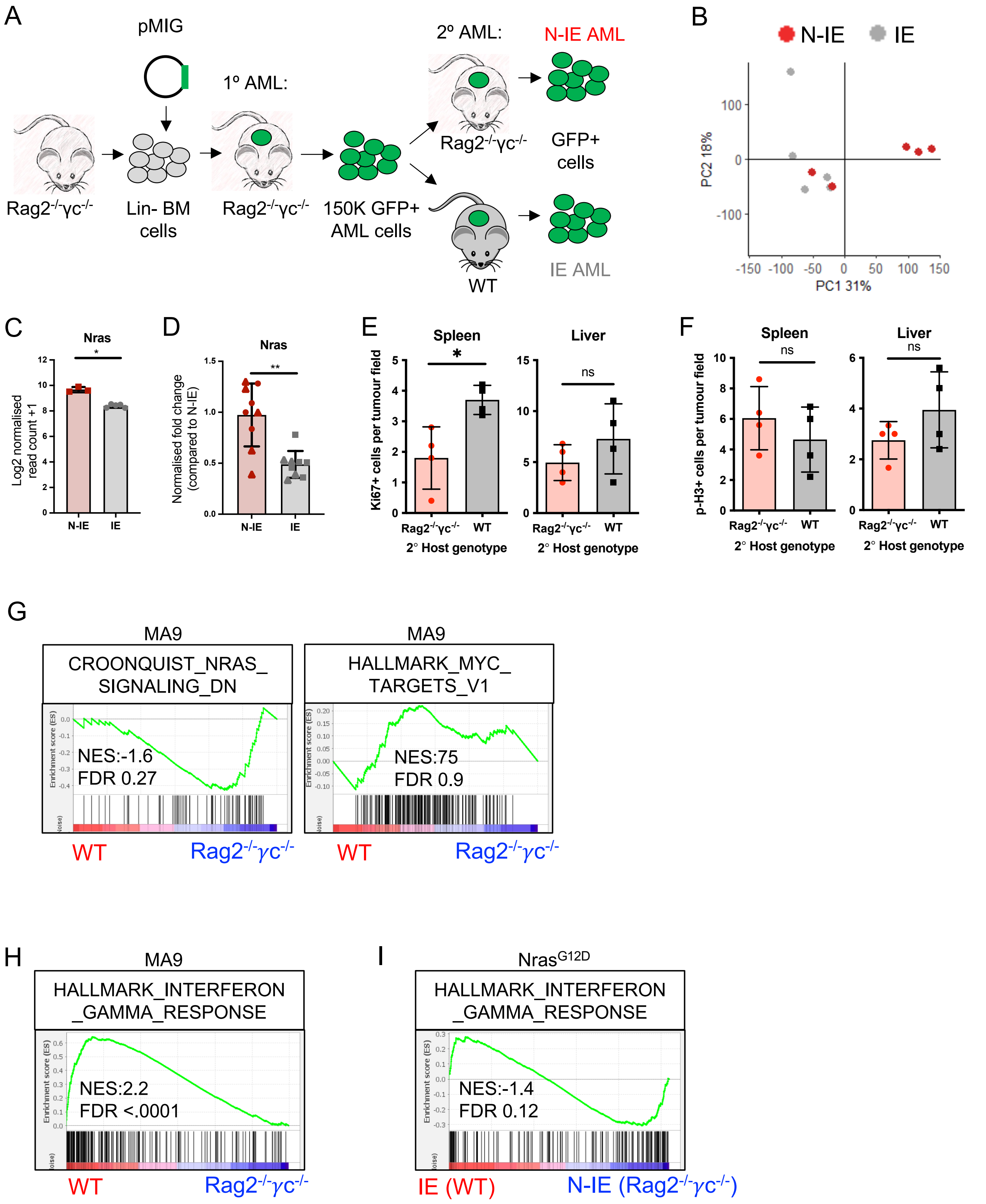


Supplementary Figure 6
